# Polymerase chain reaction-based assays facilitate the breeding and study of mouse models of Klinefelter syndrome

**DOI:** 10.1101/2021.01.02.425102

**Authors:** Haixia Zhang, Wenyan Xu, Yulin Zhou, Xiaolu Chen, Jiayang Jiang, Xiaoman Zhou, Zengge Wang, Rongqin Ke, Qiwei Guo

## Abstract

Klinefelter syndrome (KS) is one of the most frequent genetic abnormalities and the leading genetic cause of non-obstructive azoospermia. The breeding of mouse models of KS and their study are essential to advance our knowledge of the pathologic mechanism. Karyotyping and fluorescence *in situ* hybridization are reliable methods for identifying chromosomal contents. However, technical issues associated with these methods can decrease the efficiency of breeding KS mouse models and limit studies that require rapid identification of target mice. To overcome these limitations, we developed three polymerase chain reaction-based assays to measure the specific genetic information, including the presence or absence of *Sry*, copy number of *Amelx*, and Xist RNA transcript levels. Through a combined analysis of the assay results, we can infer the karyotype of target mice. We confirmed the utility of our assays with the successful generation of KS mouse models. Our assays are rapid, inexpensive, high capacity, easy to perform, and require small amounts of sample. Therefore, they facilitate the breeding and study of KS mouse models and help advance our knowledge of the pathologic mechanism underlying KS.

## Introduction

Klinefelter syndrome (KS), a set of symptoms that results from an extra X chromosome in males, is one of the most frequent genetic abnormalities and the leading genetic cause of non-obstructive azoospermia. ^1^ The prevalence of KS in newborn, infertile, and azoospermic males is approximately 0.15%, 3–4%, and 10–12%, respectively. ^2, 3^ The phenotypic spectrum of KS is wide, ranging from presenting only small testes and infertility to the classical traits and comorbidities such as hypergonadotropic hypogonadism, infertility, neurocognitive deficits, psychiatric disorders, obesity, diabetes, osteoporosis, and autoimmune disorders. ^4^ Early diagnosis of KS and subsequent treatment and intervention, such as testosterone replacement therapy, testicular sperm extraction, early speech and occupation therapy, and educational assistance, have been shown to improve the long-term quality of life for patients with KS and alleviate later complications. ^5–10^ However, the data was limited and more rigorous scientific investigations are needed. ^11^ In fact, the pathologic mechanism between abovementioned phenotypes and the extra X chromosome is much more complicated than previously envisioned and remain largely unknown, in large part because in-depth investigations, such as developmental studies and experimental manipulations, are almost impossible to perform in patients with KS for ethical reasons. ^11, 12^ Therefore, studies of animal models are essential to advance our knowledge of the pathologic mechanism of this prevalent syndrome. ^12^

The discovery of a mutant mouse line, the Y* mouse, has enabled us to generate KS mouse models. ^13^ As shown in Fig. 1, the Y* chromosome contains the X-centromere, partial pseudoautosomal region (PAR) of the X chromosome, partial PAR of the Y chromosome, and the entire non-recombining region of the Y chromosome (NRY). In addition, a small fragment of the non-pseudoautosomal region of the X chromosome (NPX), containing eight genes including *Amelx*, is located between the X-centromere and the duplicated PAR. Through cross-breeding, 40,XX^Y*^ and 41,XXY mice could be generated in the second and fourth generations, respectively. ^14^ These two mouse breeds have many of the characteristics of KS, including small firm testes, germ cell loss, hypergonadotropic hypogonadal endocrine changes, altered body proportions, behavioral and cognitive issues, and so on. ^12, 14^ Therefore, these mouse breeds have been widely used as animal models to overcome the limitations of human studies to explore the pathological mechanisms underlying KS. ^15, 16^

**Figure 1.**
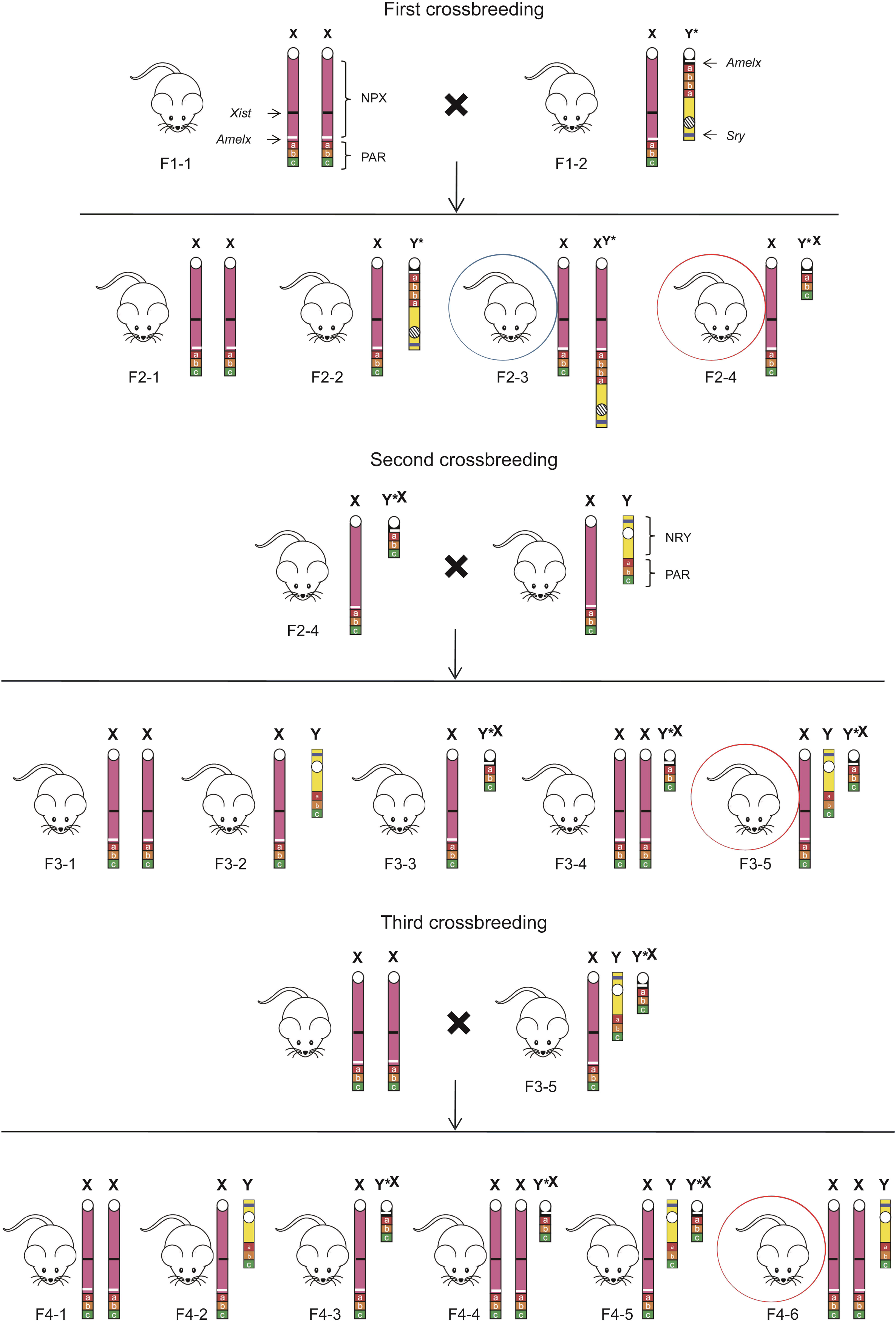
Breeding of Klinefelter syndrome (KS) mouse models. The mouse in the blue circle is a 40,XX^Y*^ mouse, i.e., one of the KS mouse models. The mice in the red circles are the target mice in each generation for breeding 41,XXY mice, i.e., another KS mouse model. This figure was modified from figures in a previous study (Front Neuroendocrinol 2014; 35: 405-19). NPX: non-pseudoautosomal region of the X chromosome; NRY: non-recombining region of the Y chromosome; PAR: pseudoautosomal region.

One of the most important steps in the successful breeding of 40,XX^Y*^ and 41,XXY mice is the accurate identification of the karyotypes and chromosomal fragments of the targeted mouse breeds (Fig. 1). This is typically performed through cytogenetic methods, such as karyotyping or fluorescence *in situ* hybridization (FISH). To ensure that the mice are alive for subsequent breeding after testing, culture cells derived from blood samples or tissue biopsies are used. ^17^ However, identification of target mice by karyotyping or FISH is labor-intensive, time-consuming (5–10 days), and requires a relatively high level of expertise. These drawbacks reduce their usefulness of large-scale detection, which is generally required to generate enough model mice for study use. Moreover, the pathological changes in the KS model mice, such as germ cell loss, occur prenatally and progress rapidly after birth. ^15, 18^ Therefore, studies of these early pathological changes require immediate identification of target mice prenatally or neonatally; this poses a challenge for karyotyping and FISH, which require a relatively long turnaround time.

To overcome these limitations, we developed three polymerase chain reaction (PCR)-based assays to identify the chromosomal contents of target mice and confirmed the utility of our assays for reliable breeding of KS mouse models.

## Materials and Methods

### Study design

As shown in Fig. 1, each karyotype has specific sex chromosome contents that could provide specific genetic information such as gene copy number and transcriptional activity. Theoretically, by analyzing this genetic information using molecular methods, instead of cytogenetic analyses such as karyotyping and FISH, we would be able to infer the karyotype of a mouse breed. Based on this concept, three genes, *Sry, Amelx*, and *Xist*, could be informative. *Sry* is located in the NRY, and its presence or absence determines the gender of the mouse. Since *Amelx* is located in the NPX and is also present on the Y* chromosome, the copy number of *Amelx* implies the copy number of the NPX fragment. ^19^ *Xist* is also located on the NPX and is transcribed into a long noncoding RNA that can transcriptionally silence one of the two X chromosomes to achieve dosage equivalence between males and females. Thus, a high level of Xist RNA is expected when there are two X chromosomes in a single cell, whereas an absence or low level of Xist RNA is expected when there is only one X chromosome in a single cell. ^20^ Corresponding to the breeding protocol shown in Fig. 1, the genetic information of the mouse breeds in each generation is shown in Fig. 2. This genetic information was determined using the PCR-based assays designed to measure the presence or absence of *Sry*, copy number of *Amelx*, and the transcript levels of Xist RNA. Notably, the combined information for two genes was sufficient to identify a target karyotype (Fig. 2).

**Figure 2.**
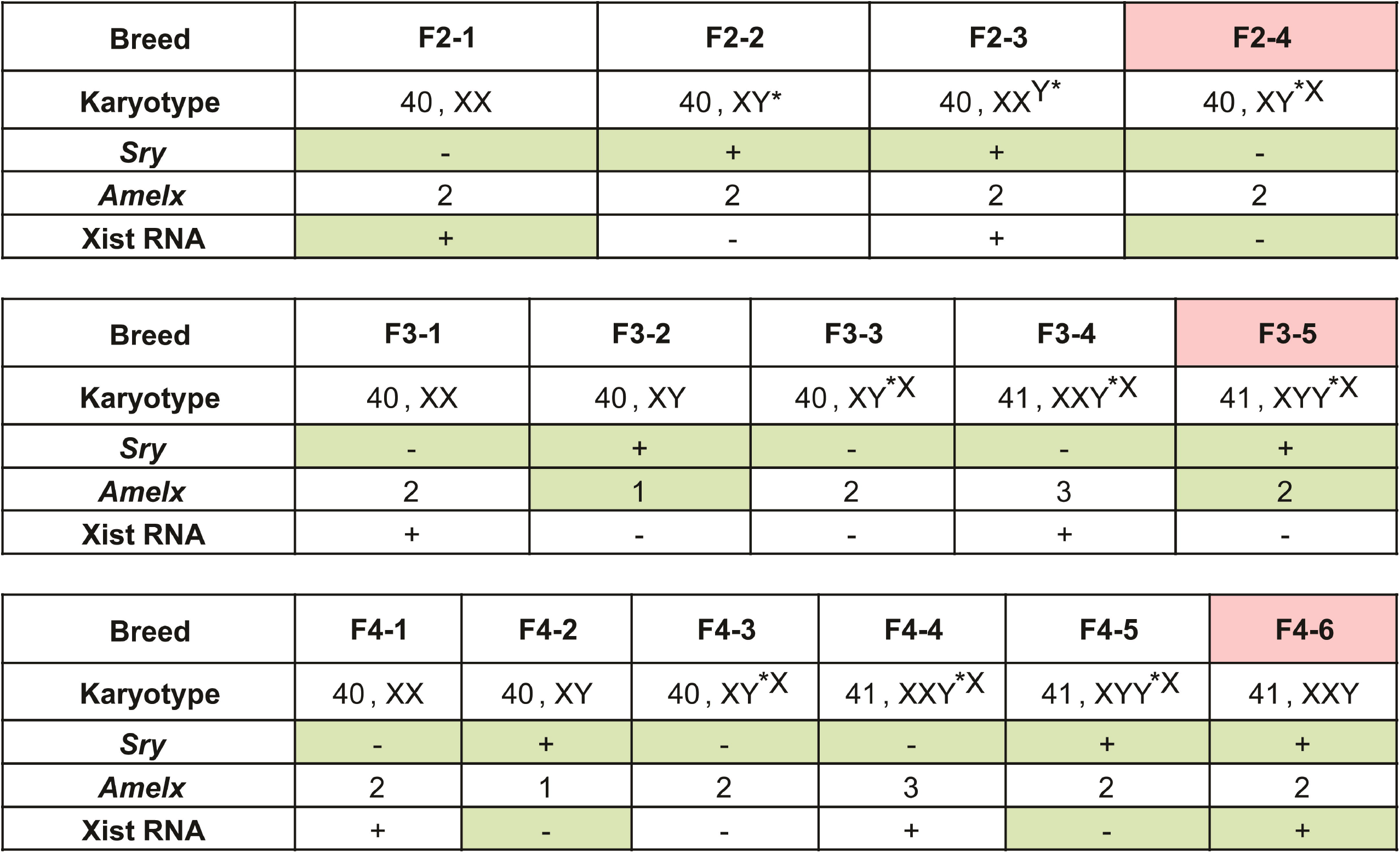
Genetic information of the mouse breeds obtained during the breeding of Klinefelter syndrome mouse models. +: presence of *Sry* or a high Xist RNA transcript level; –: absence of *Sry* or low/no Xist RNA transcript. Pink labels indicate the target mice in each generation for the breeding of 41,XXY mice. Green labels indicate the assays required to identify the target mice in each generation. For example, in the second generation, each mouse was first tested for *Sry*, and those negative for *Sry* were further tested for Xist RNA transcript levels. Mice with low/no Xist RNA transcript, i.e., negative results in both assays, were identified as 40,XY*^X^ mice.

### Mice and samples

To establish the PCR assays, tail tissue samples from 16 pairs of 40,XX and 40,XY mice (C57BL/6J) were obtained from the Experimental Teaching Department, School of Medicine, Xiamen University. To generate 40,XX^Y*^ and 41,XXY mice, breeding pairs of 40,XY* (C57BL/6JEiJ) and 40,XX (C57BL/6J) mice were purchased from The Jackson Laboratory (Bar Harbor, ME, USA). The 40,XY mice (C57BL/6J) were purchased from Xiamen University Laboratory Animal Center. The breeding was performed in the standard animal facility of the Xiamen University Laboratory Animal Center.

### Nucleic acid purification

Approximately 20 mg of tail tissue was used to purify DNA or RNA with the QIAamp Fast DNA Tissue Kit (Qiagen, Valencia, CA, USA) or the RNeasy^®^ mini Kit (Qiagen), respectively, according to the manufacturer’s protocols. The purity and concentration of the purified nucleic acid were determined by measuring the absorbance at 260 nm and 280 nm using a NanoDrop 2000 spectrophotometer (Thermo Fisher, Waltham, MA, USA).

### Reaction conditions

The PCR conditions for all assays were identical: each 25 μL reaction contained 10 mmol/L Tris-HCl (pH 8.3), 50 mmol/L KCl, 1 U of TaqHS (Takara, Dalian, China), 2 mmol/L Mg^2+^, 0.2 mmol/L each dNTP, 0.8× LightCycler 480 ResoLight Dye (Roche Applied Science GmbH, Mannheim, Germany), 0.2 mmol/L each forward and reverse primer, and a specific amount of DNA or cDNA template. The amplification cycling conditions were as follows: 95 °C for 3 min, followed by 40 cycles of 95 °C for 20 s, 60 °C for 20 s, and 72 °C for 20 s. Fluorescence was recorded at the end of each annealing step. The sequences of the primers used in each assay are listed in Table 1. For the measurement of *Sry*, a melting analysis was performed after the amplification, which began with denaturation at 95 °C for 1 min and renaturation at 35 °C for 3 min, followed by melting from 40 °C to 95 °C, with a ramp rate of 0.04 °C/s. Fluorescence was recorded every 0.3 °C. PCR and melting analysis were performed on a SLAN-96S thermocycler (Zeesan, Xiamen, China). Before PCR, RNA samples were reverse transcribed with the GoScript Reverse Transcription System (Promega, Beijing, China) according to the manufacturer’s protocol, and 1 μL of the resulting cDNA was used as template for subsequent PCR.

**Table 1.**
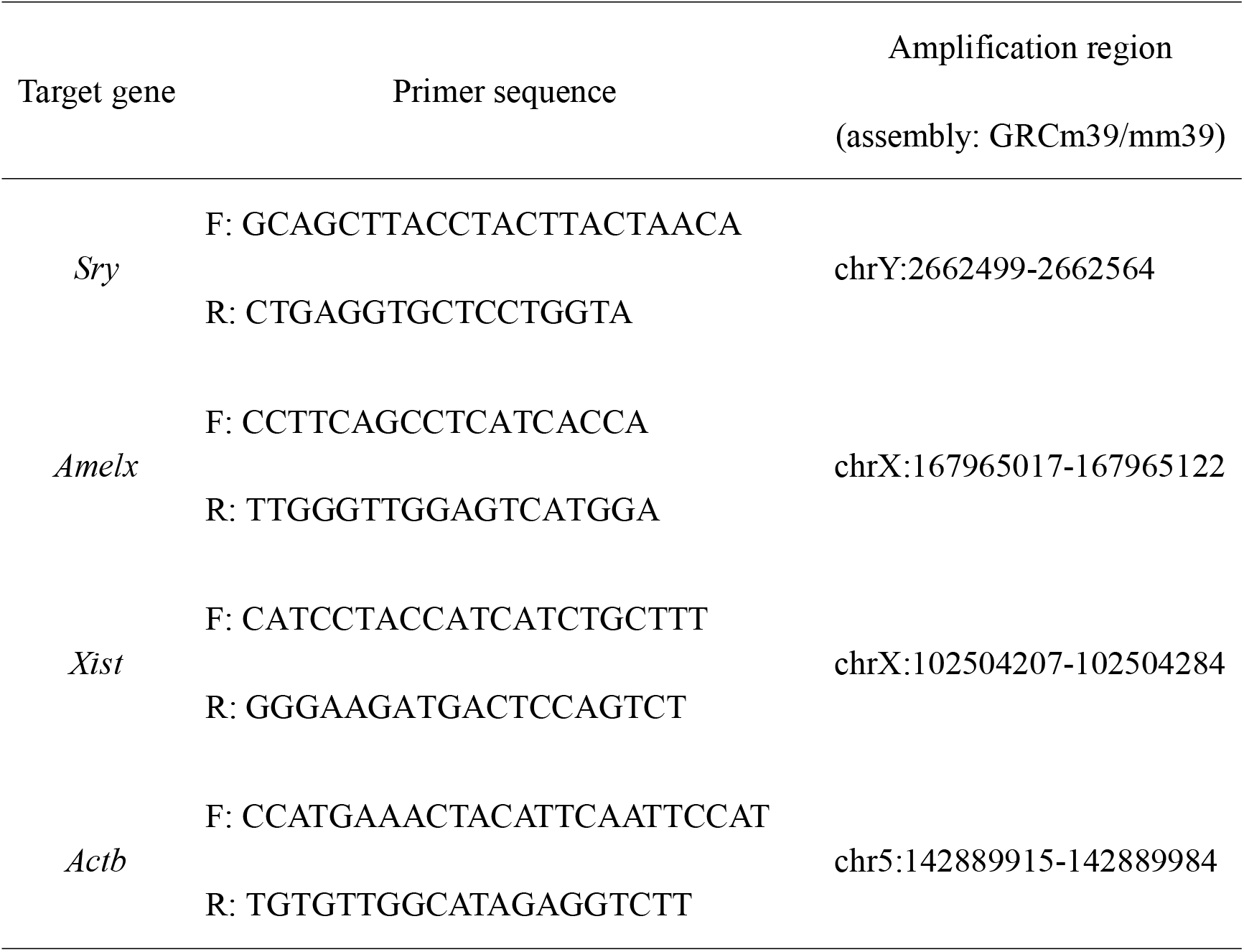
Primer information of PCR-based assays.

### Data analysis

For the analysis of *Sry*, a melting profile was used to determine its presence or absence. To quantify the copy number of *Amelx*, relative quantification was performed by measuring the difference between the quantification cycle (Cq) values obtained for *Amelx* and *Actb* (ΔCq_Amelx - Actb_). Similarly, to evaluate Xist RNA transcript levels, relative quantification was performed by measuring the difference between the Cq values for the *Xist* and *Actb* transcripts (ΔCq_Xist - Actb_). *Actb* was used as a quantitative reference because it has consistently has two copies and a stable transcriptional pattern in mouse cells. The Cq value is defined as the amplification cycle when the fluorescence intensity of an amplicon reaches a specific threshold.^21^ Based on a previous study, the threshold was set to 30% of the plateau fluorescence intensity as the optimal Cq value for differentiation of different karyotypes. ^22^

### Karyotyping and histologic analysis

To confirm the accuracy of our assays, after breeding, the karyotypes of the target mice were examined by karyotyping using hemopoietic lineage cells derived from the bone marrow according to a previous protocol. ^17^ For histologic analysis of the testes of KS model mice, periodic acid-Schiff staining was performed on cross-sections of the seminiferous tubules according to a previous protocol. ^23^

### Ethics statement

Mouse breeding and experimentation were performed in accordance with international, national, and institutional guidelines for the ethical use of animals. ^24^ The study was approved by the Ethics Committee of Xiamen University Laboratory Animal Center.

## Results

### Establishing an assay for the detection of *Sry*

To determine the optimal amount of DNA template for the *Sry* detection assay, serial dilutions of DNA isolated from 40,XY (*Sry* positive) and 40,XX (*Sry* negative) mice were evaluated. A no template control was also examined to determine the melting profiles of primer dimers. As shown in Fig. 3A, the melting profiles changed with the amount of DNA template. To obtain only the *Sry* amplicon, at least 2 ng of DNA template was required for samples from the 40,XY mouse. However, for the 40,XX mouse samples, the use of a larger amount of DNA template (≥20 ng) generated non-specific amplicons that presented similar melting profiles to those of the *Sry* amplicons. Therefore, 2 ng was considered as the optimal amount of template to specifically detect the presence or absence of *Sry* and was used for subsequent experiments.

**Figure 3.**
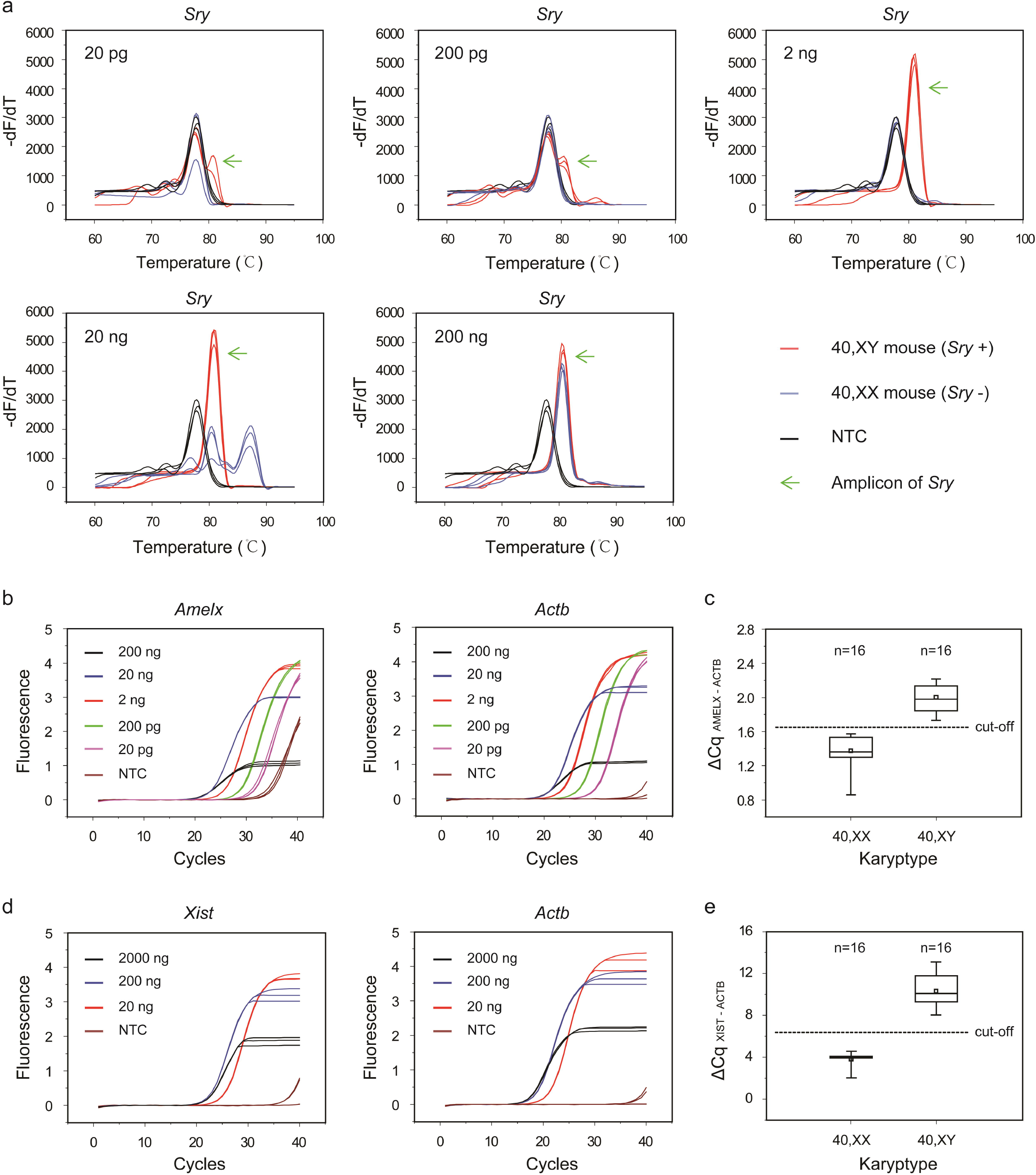
PCR-based assays for the breeding of Klinefelter syndrome mouse models. (a) Melting profiles for detecting *Sry* using different amounts of DNA template. (b) Assays to quantify *Amelx* using different amounts of DNA template. (c) Establishing a cut-off value for *Amelx* quantification. (d) Assays for evaluating Xist RNA transcript levels using different amounts of RNA template. (e) Establishing a cut-off value for Xist RNA quantification. Each sample was tested in triplicate, and the Cq value of each sample is the mean Cq value of three replicates. NTC: no template control.

### Establishing an assay for quantification of *Amelx*

We first evaluated the influence of the amount of DNA template on the effectiveness of the *Amelx* quantification assay by examining serial dilutions of DNA isolated from a 40,XX mouse as a template. As shown in Fig. 3B, amplification in both the *Amelx* and *Actb* detection reactions was robust across a wide range of template concentrations (20 pg–200 ng). However, a repressive effect was observed when a relatively large amount of DNA template (≥20 ng) was used. Therefore, 2 ng was chosen as the amount of template to evaluate the ranges of the ΔCq_Amelx - Actb_ values for the 40,XX (two copies of *Amelx*) and 40,XY (one copy of *Amelx*) mouse samples. As shown in Fig. 3C, the ranges for 16 samples from 40,XX and 40,XY mice were distinct from each other. The mean ΔCq_Amelx - Actb_ value (1.65) of the maximum of the 40,XX samples and the minimum of the 40,XY samples was used as the cut-off value for classifying mice as having one or two copies of *Amelx* in subsequent experiments.

### Establishing an assay for evaluating Xist RNA transcript levels

We first evaluated the influence of different amounts (2000, 200, and 20 ng) of RNA template from a 40,XX mouse (high level of Xist RNA transcripts) in a reverse transcription reaction on PCR effectiveness. As shown in Fig. 3D, both 2 000 ng and 200 ng of RNA had a repressive effect. Therefore, 20 ng of RNA template was used for reverse transcription to evaluate the range of the ΔCq_Xist - Actb_ values for 40,XX and 40,XY mouse samples (no or low level of Xist RNA transcript). As shown in Fig. 3E, the range of the ΔCq_Xist - Actb_ values for the 16 40,XX and 40,XY mouse samples were distinct from each other. The mean ΔCq_Xist - Actb_ value (6.05) of the maximum for the 40,XX samples and the minimum for the 40,XY samples was used as the cut-off value for differentiating Xist RNA transcript levels in subsequent experiments.

### Breeding KS mouse models using these PCR assays

Using the established PCR assays, we bred 40,XX^Y*^ and 41,XXY mice from a pair of 40, XY* and 40,XX mice following the protocol in Fig. 1, which confirmed the effectiveness of our method. Ultimately, we obtained one 40,XX^Y*^ mouse and four 41,XXY mice. Subsequent karyotyping and histological analysis confirmed the reliability of our assays (Figs. 4 and 5).

**Figure 4.**
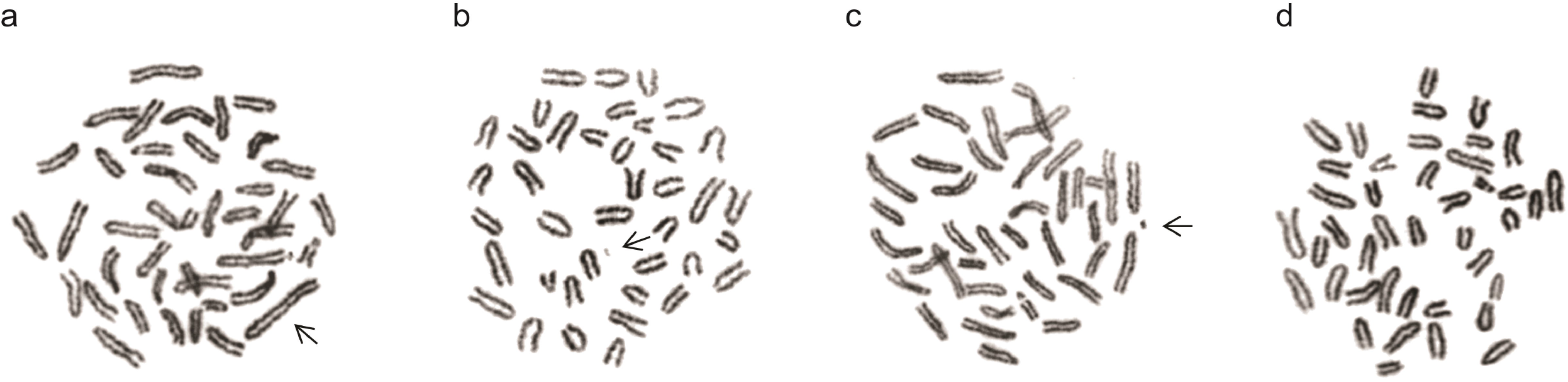
Karyotypes of the target mice for breeding Klinefelter syndrome mouse models. (a) A 40,XX^Y*^ mouse from the second generation. The arrow indicates the X^Y*^ chromosome. (b) A 40,XY^X*^ mouse from the second generation. The arrow indicates the Y^X*^ chromosome. (c) A 41,XYY^X*^ mouse from the third generation. The arrow indicates the Yx* chromosome. (d) A 41,XXY mouse from the fourth generation.

**Figure 5.**
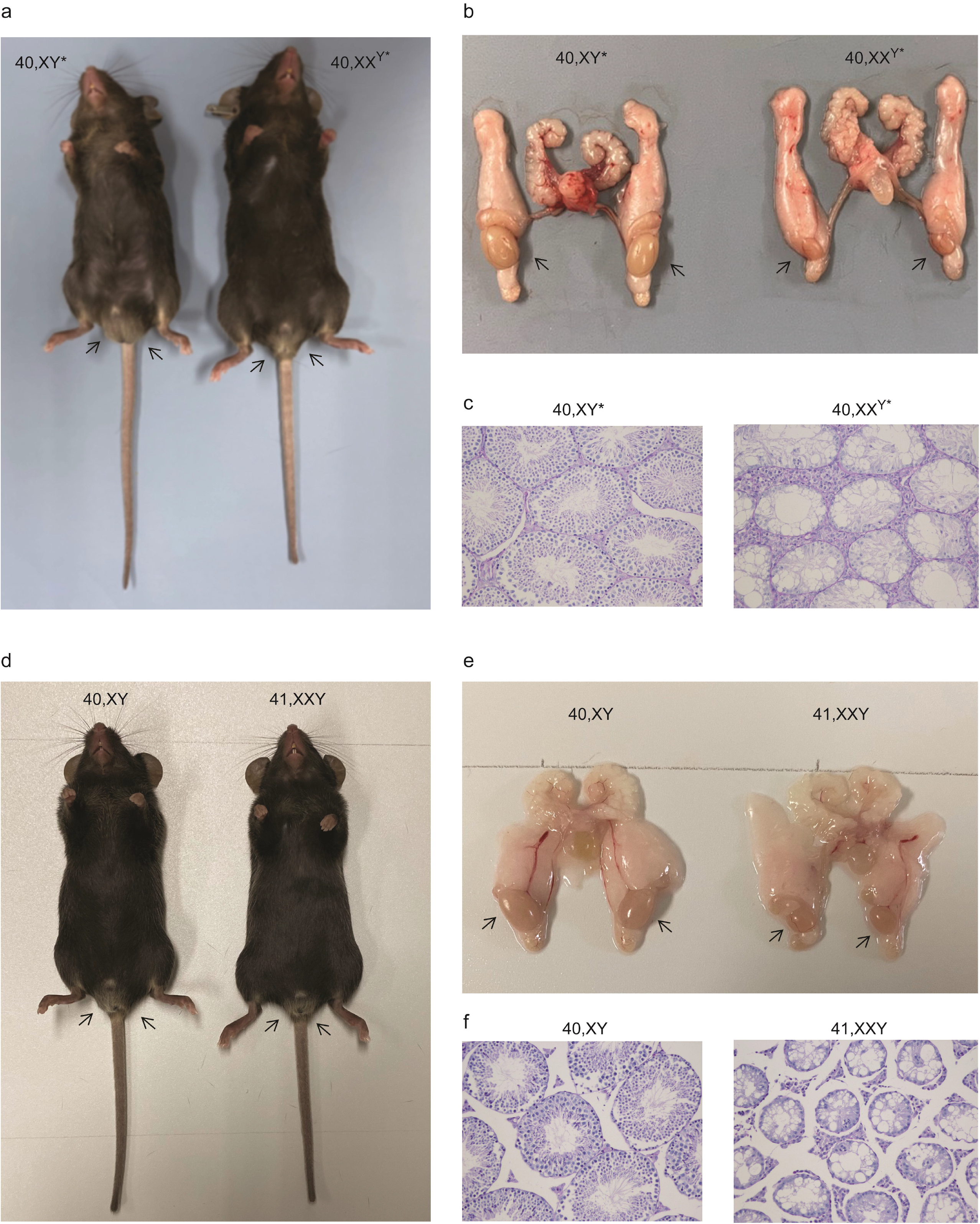
Testicular phenotypes of Klinefelter syndrome mouse models. (a-c) Respective appearance, anatomic analysis, and histologic analysis of the testes of a 40,XX^Y*^ mouse and 40,X^Y*^ littermate. (d-f) Respective appearance, anatomic analysis, and histologic analysis of the testes of a 41,XXY mouse and 40,XY littermate. The arrows indicate the position of the testes. As expected, the 40,XX^Y*^ and 41,XXY mice presented small, firm testes. Moreover, the tubule diameters of 40,X^Y*^ and 41,XXY mice were smaller than those of their reference littermates, germ cells were absent, and only Sertoli cells were observed within the tubules.

## Discussion

Karyotyping and FISH are reliable methods that have been used for decades to identify chromosomal abnormalities, which are essential for the breeding and study of KS mouse models. However, these methods have some technical issues, e.g., a relatively low capacity and long turnaround time, that decrease the breeding efficiency and limit studies which require rapid identification of chromosomal contents.

PCR is a basic tool of the molecular laboratory that has demonstrated capability for detecting various types of genetic abnormalities through different data analysis strategies. For example, we can detect the presence or absence of a specific gene region (e.g., *AZF* on the Y chromosome) by analyzing amplification profiles, ^25^ and we can detect numerical changes (e.g., trisomy 21, 47,XXY, or a *SMA* carrier) by analyzing meting profiles or ΔCq values. ^5, 22, 26–28^ Based on previous studies, we developed three PCR-based assays to detect genetic information related to KS, including the presence or absence of *Sry*, changes in the copy number of *Amelx*, and Xist RNA transcript levels in mouse tissue samples. Using these assays, we were able to identify target mice for the breeding of KS models without performing karyotyping or FISH (Figs. 1 and 2). Compared to karyotyping or FISH, our method is rapid, as it takes less than 1 h for nucleic acid purification, less than 2 h for reverse transcription, and less than 2 h for PCR and melting analysis. A short turnaround time for mouse identification is essential for studies focused on rapid physical and pathologic changes in fetal or newborn mice, e.g., testicular degeneration. ^15, 18^ Our method is also inexpensive (<0.5 US dollar per PCR assay), high capacity (up to 96 samples per assay), and easy to perform. Moreover, only 2 ng of DNA and 20 ng of RNA was sufficient to obtain reliable results (Fig. 3), suggesting that less than 0.2 mg of tissue or 1 μl of peripheral blood would be a sufficient sample, further revealing its utility, especially for the detection of fetal or newborn mice.

An ideal assay for *Sry* identification should only yield a product from male mouse samples with *Sry* but not from female mouse samples without *Sry*, so that we can identify male mouse samples directly by their amplification profiles. In practice, however, non-specific amplicons derived from paralogous sequences and/or primer dimers were difficult to eliminate from reactions with female mouse samples. Because of this, melting profiles are more specific than the amplification profiles since the melting profiles of the *Sry* amplicons could be readily differentiated from those of non-specific amplicons (Fig. 3A). The amount of DNA template used plays an important role in the melting profiles, and 2 ng was confirmed to be an optimal amount of template in our study (Fig. 3A). However, if different PCR reagents are used, the optimal amount of DNA template should be re-evaluated.

Theoretically, after equalizing the DNA template input for the *Amelx* assay, samples with different *Amelx* copy numbers should generate different ranges of Cq values. Therefore, we could infer the *Amelx* copy number in a mouse sample by its Cq value. In practice, we confirmed this assumption (data not shown). However, to further diminish the potential impacts of slight differences (e.g., differences in the quantity or quality of the DNA template) between samples, *Actb* was used as a reference gene to calibrate the *Amelx* quantification. Accordingly, instead of ranges of Cq values, ranges of ΔCq_Amelx - Actb_ values were established to quantify *Amelx* in unknown samples (Fig. 3C). Moreover, rather than using confidence intervals for the different ranges of ΔCq_Amelx - Actb_ values, we used a single cut-off value to determine the *Amelx* copy number (Fig. 3C), which simplified the data analysis.

The underlying principle of the assay used to determine Xist RNA transcript levels was identical to that of *Amelx* quantification. Similarly, a large amount of nucleic acid template had a repressive effect on PCR amplification (Fig. 3D). We further evaluated whether the repressive effect was derived from the large amount of RNA or the resulting cDNA. The results suggested that the repressive effect was greater for large amounts of RNA (data not shown). Therefore, a smaller amount of RNA was preferable for reverse transcription, which was sufficient to reliably determine Xist RNA transcript levels (Fig. 3E).

In conclusion, we developed three PCR-based assays for facilitating the breeding and study of KS mouse models. Our method is rapid, inexpensive, high capacity, easy to perform, and requires small amounts of sample. We confirmed the utility of our assays for the successful generation of KS mouse models. We believe our method will advance the knowledge of the pathologic mechanism of KS.

## Authors’ Contributions

HXZ, WYX and YLZ participated in study design and the establishment of PCR assays. XLC, JYJ, XMZ, ZGW, and RQK participated in breeding of KS mouse models. QWG conceived of the study, and participated in its design and coordination and draft the manuscript. All authors have read and approved the final version of the manuscript, and agree with the order of presentation of the authors.

## Acknowledgements

We thank the Experimental Teaching Department, School of Medicine, Xiamen University and the Xiamen University Laboratory Animal Center for their kind supports to this study.

## Conflict of Interest

None of the authors declare competing financial interests.

## Notes

### Competing Interest Statement

The authors have declared no competing interest.

